# Structure and flexibility of the DNA polymerase holoenzyme of vaccinia virus

**DOI:** 10.1101/2023.09.03.556150

**Authors:** Wim P. Burmeister, Laetitia Boutin, Aurelia C. Balestra, Henri Gröger, Allison Ballandras-Colas, Stephanie Hutin, Christian Kraft, Clemens Grimm, Bettina Böttcher, Utz Fischer, Nicolas Tarbouriech, Frédéric Iseni

**Affiliations:** Institut de Biologie Structurale (IBS), Université Grenoble Alpes (UGA), Commissariat à l’Energie Atomique et aux Energies Alternatives (CEA), Centre National de la Recherche Scientifique (CNRS), Grenoble, France; Institut de Recherche Biomédicale des Armées, Brétigny-sur-Orge, France; Biozentrum, Universität Würzburg, Würzburg, Germany

## Abstract

The year 2022 was marked by the mpox outbreak caused by human monkeypox virus (MPXV), which is about 98 % identical to vaccinia virus (VACV) at the sequence level regarding the proteins involved in DNA replication. We present the strategy for the production of the VACV DNA polymerase holoenzyme composed of the E9 polymerase associated with its co-factor, the A20-D4 heterodimer, which led to the 3.8 Å cryo-electron microscopy (cryo-EM) structure of the DNA-free form of the holoenzyme. Model building used high-resolution structures of components of the complex and the A20 structure predicted by AlphaFold 2. The structure of E9 does not change in context of the holoenzyme compared to the crystal structure. As for the MPXV holoenzyme, a contact between E9 and D4 is mediated by a cluster of hydrophobic residues. The holoenzyme structure is quite compact and surprisingly similar to the MPXV holoenzyme in presence of a DNA template, with the exception of a movement of the finger domain and the thumb domain, which becomes ordered in presence of DNA. Even in absence of DNA, the VACV holoenzyme structure is too compact for an agreement with SAXS data. This suggests the presence of more open conformations in solution, which are also predicted by Alphafold 2 indicating hinge regions located within A20. Using biolayer interferometry we showed that indeed, the E9-D4 interaction is weak and transient although very important as it has not been possible to obtain viable viruses carrying mutations of key residues in the E9-D4 interface.

**Author Summary:** The 2022 outbreak of mpox is caused by monkeypox virus closely related to the best studied model, vaccinia virus. Genome replication, which takes place largely autonomously in the cytosol of the infected cell, is still not really understood. Viral DNA synthesis involves a DNA repair enzyme, the uracil-DNA glycosylase D4 linked to the structural protein A20 forming the processivity factor, which in turn binds to E9 forming the complex required for processive DNA synthesis. Here we present the first structure of the vaccinia virus polymerase holoenzyme E9-A20-D4 at 3.8 Å obtained by cryo-electron microscopy. This structure, together with several recent structures from monkeypox virus, provide a static view of the complex with a previously undescribed contact between E9 and D4. Our small-angle scattering data show that other conformations, taking advantage of 2 hinge regions in A20, exist in solution. Using site-directed mutagenesis and binding studies we show that the contact between E9 and D4, which serves to encircle the template strand, is important, but transient. Thus the current model of the orientation of the holoenzyme on the replication fork may not be the only one possible.

## Introduction

With the 2022 epidemic outbreak of mpox caused by human monkeypox virus (MPXV) of clade IIb, poxviruses are again in the headlines. Previous human monkeypox infections mainly remained localized in West and Central Africa, where viruses from clades I and IIa circulate in rodent reservoirs [1]. In humans, the two clades lead to zoonotic infections with different degrees of severity. In the past, sporadic infections occurred outside Africa and have always been transmitted by pet rodents. For the first time, the 2022 outbreak involved human to human transmission outside Africa leading to an acceleration of the evolution of the viral genome. Human monkeypox virus is mainly transmitted by sexual contacts between men with characteristic skin lesions in the oral and ano-genital area, fever and myalgia, which could be followed by centrifugal secondary eruptions [2]. Available antivirals are tecovirimat [3] interfering with virion assembly and brincidofovir, which targets viral DNA replication through an action as chain terminator (see [4] for a review). MPXV is closely related to vaccinia virus (VACV), the best-studied orthopoxvirus, initially used for smallpox eradication by vaccination. It replicates and assembles in perinuclear viral factories where viral DNA synthesis takes place independently of the host cell nucleus [5]. To date, numerous aspects of the replication cycle remain unsolved. The VACV genome is a 196 kbp double-stranded DNA circularized at the extremities with peculiar telomere structures with imperfect base pairing preceded by repeat sequences [5]. It has been proposed that an origin of replication is located within the telomere [6]. The replication mechanism in still controversial and proceeds most likely through a variant of a rolling circle mechanism suggested by the presence of a primase activity [7] and Okazaki fragments [8]. In addition to the polymerase holoenzyme, the helicase-primase D5 and the single-stranded DNA binding protein I3 are required for DNA replication [5]. There has been a long quest for the structure of the poxvirus DNA polymerase holoenzyme since it has been shown that processive DNA synthesis by VACV E9 requires a cofactor composed of VACV A20 and D4 [9], D4 being the viral uracil-DNA glycosylase (UNG)[10]. The polymerase E9 is a member of the DNA polymerase family B possessing DNA polymerase and 3’-5’ proofreading exonuclease activities [11]. Interestingly, it was also shown to catalyze annealing of single-stranded DNA [12], an activity not found in other family B DNA polymerases. The end-joining reaction requires the 3’-5’ exonuclease activity of E9 that degrades the extremities of dsDNA to create 5’-ssDNA overhangs. E9 on its own was shown to be distributive under physiological conditions, adding only few nucleotides per binding event [13]. An initial low resolution model of the complex [14] has been completed progressively by high-resolution structural information on several proteins and subcomplexes [15–17]. Still, the mechanism of the processivity factor remained largely unknown. Sparked by the mpox outbreak, interest switched from VACV to MPXV where the proteins of the DNA replication machinery are about 98 % identical at the amino acid sequence level. The VACV nomenclature of the reading frames is used throughout the article although the E9-A20-D4 complex is named F8-A22-E4 for MPXV. A first high-resolution structure of the polymerase holoenzyme with bound template DNA, primer strand and incoming dTTP nucleotide obtained by cryo-electron microscopy (cryo-EM) was published by Peng and coworkers [18]. It showed a previously unidentified contact between E9 and D4, which allows the E9-A20-D4 complex to encircle the template strand leading to an unexpectedly compact structure of the holoenzyme. In this model, A20 appears to play just a role of a connector and the active site of D4 with its DNA-binding capacity [19] is not used.

Further structures of the MPXV polymerase holoenzyme [20,21] showed a DNA-free state with a similar circular structure and an E9-D4 contact, but also a dimer of trimers [20].

We present a system for the production of the VACV E9-A20-D4 holoenzyme and its 3.8 Å structure obtained by single particle cryo-EM. As our model shows discrepancies with data obtained in solution by small-angle X-ray scattering (SAXS) we predicted the existence of open forms of the complex leading to studies of the importance of the E9-D4 contact and a direct comparison with recently determined MPXV polymerase holoenzyme structures.

## Results

The E9-A20-D4 holoenzyme could be expressed with a good yield in insect cells using a single recombinant baculovirus. The double tag, a 6His-tag on E9 and a Strep-tag on D4, allowed an efficient purification and led to a stoichiometric complex (Fig 1AB). SEC-MALS analysis of the purified complex yielded a single peak of the complex with a mass of 185 kDa compared to a theoretical mass of 197 kDa (Fig 1C). The efficiency of the purification step on the streptactin column allowed to simplify the purification protocol for cryo-EM sample preparation by the omission of the TEV cleavage and second Ni-column purification step. The analysis of the E9-A20-D4 complex by cryo-electron microscopy was hampered by strong preferential orientations of the particles at the ice-air interface. One dataset showed less prominent preferential orientations, which allowed the 3D reconstruction of the complex (S1 Fig) and refinement to a resolution of 3.8 Å (Fig 2A). Surprisingly, the structure showed a conformation, where the processivity factor A20-D4 was folded back onto the polymerase subunit creating the D4-E9 contact described for the MPXV holoenzyme in presence of DNA [18](Fig 2B). The parts of the complex where previous high-resolution crystal structures were available were very well defined and allowed to fit E9 [16], the complex of D4 and A201-50 [15] and A20_304-426_ using its modelled position relative to E9 [17]. The least well defined part of the model of A20 corresponds to an inactive DNA ligase domain (res. 67-310) composed more precisely of a catalytic subdomain followed by an OB-domain. The ligase domain has been identified in the MPXV processivity factor protein A22 [18] and in the Alphafold 2 predicted protein structures of orthopoxviruses [22]. The Alphafold 2 prediction of A20 could be fitted reliably into the electron density adjusting the relative orientation of the domains A20_1-50_ (Fig 2c), ligase domain and A20_304-426_ although for the final structure a model based on the very similar MPXV A22 structure has been used. The flexibility around the hinge regions creates probably a disorder of the ligase domain, which limited the number of exploitable particles for 3D reconstruction and refinement (S1 Fig). The thumb domain (res. 830-1006) of E9 is not visible in the electron density showing its flexibility. In the refined model of the VACV polymerase holoenzyme, the carbon-alpha positions of the ligase domain superpose with 0.56 Å rms onto the MPXV A22 structure and with 1.13 Å rms onto the Alphafold 2 model.

**Fig 1.**
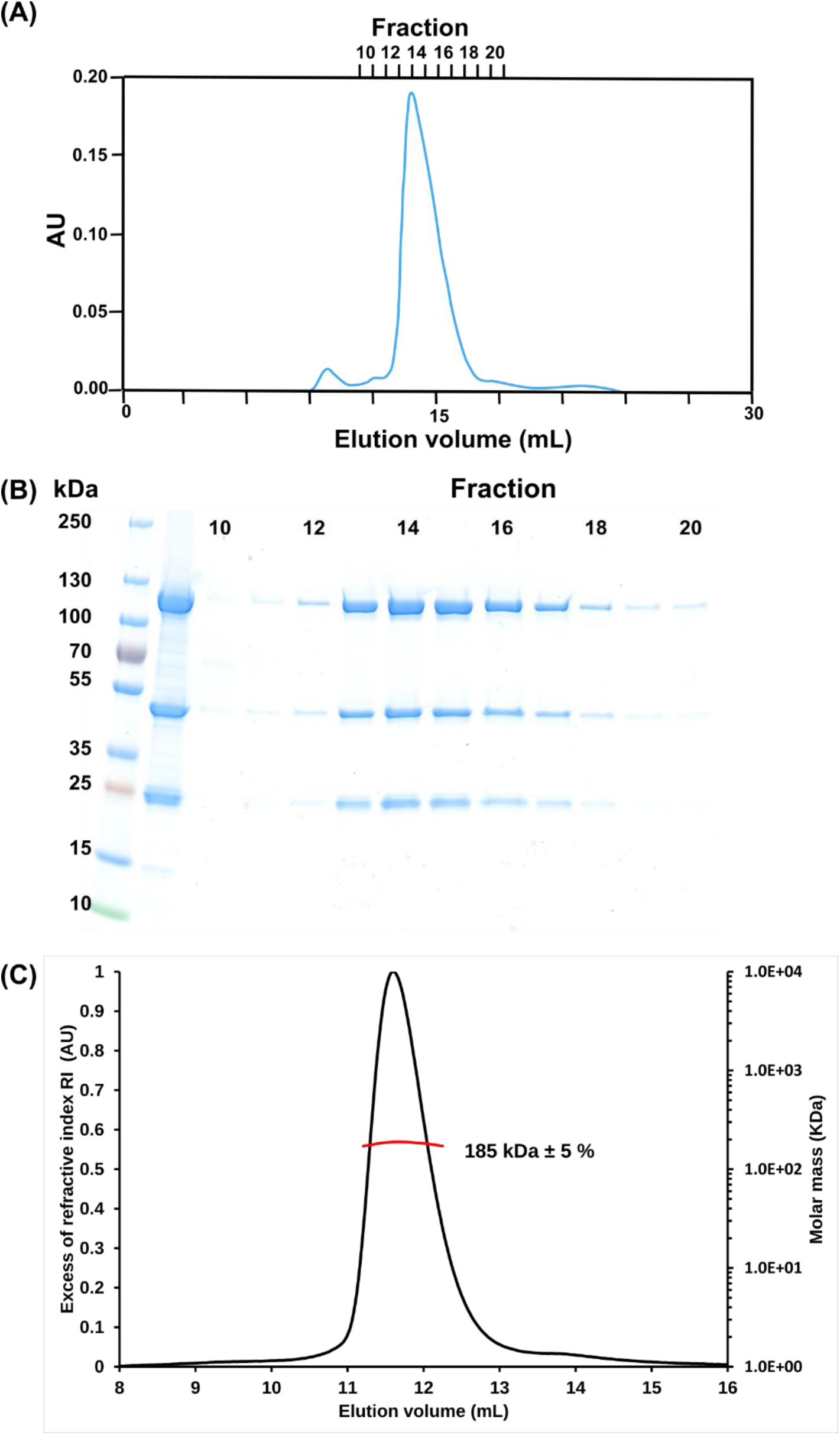
Production of the E9-A20-D4 complex. (**A**) Chromatogram of the final size exclusion step of the purification. (**B**) Fractions of panel (A) analyzed by SDS PAGE. (**C)** Result of a SEC-MALS run of the purified complex.

**Fig 2.**
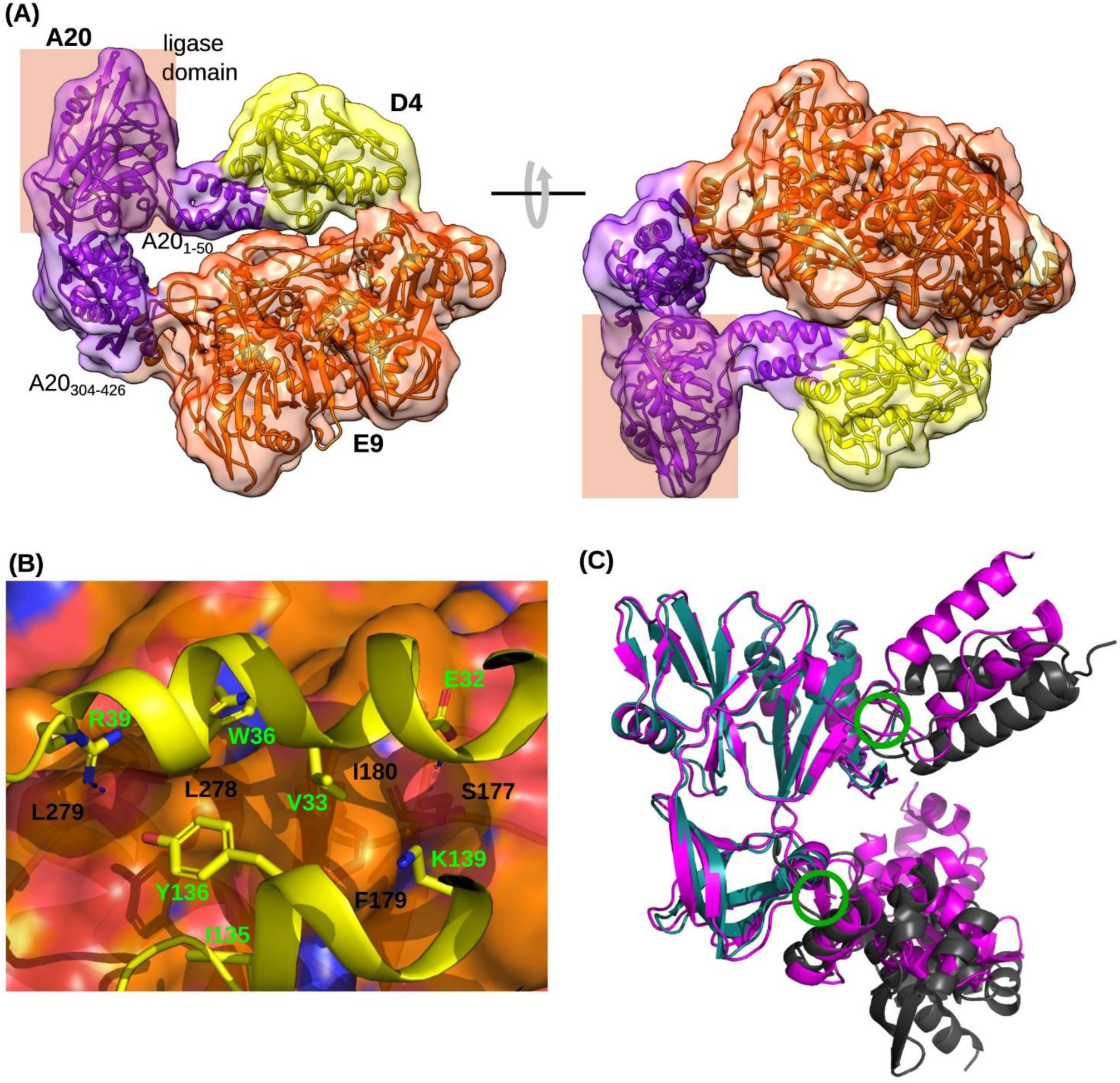
E9-A20-D4 holoenzyme structure. (**A**) The refined 3.8 Å structure of the holoenzyme is shown in cartoon representation in two views together with the electron density map before sharpening. The disordered thumb domain of E9 (res. 830 – 1006) is not shown. (**B**) The D4 binding site of E9 (orange carbon atoms) is shown in a transparent surface representation; the underlying contact residues are labelled and shown with brown carbon atoms and contact residues of D4 in a stick model (yellow carbon atoms). (**C**) Superposition of A20 in the context of the VACV polymerase holoenzyme heterotrimer (violet) with the Alphafold 2 prediction (grey). Green circles indicate hinges.

The E9-D4 contact involves mainly two α-helices of D4 contributing E32, V33, W36 and R39, I135, Y136 and K139 while on E9 it involves S177, F179, I180, L278 and L279 (Fig 2B). The side chains of D4 E32 and E9 S177, as well as D4 R39 and the main chain carbonyl of E9 L278 could be involved in hydrogen bonds. This interface belongs to one of the best defined parts of the complex with a local resolution below 4 Å (S1E Fig). The above mentioned residues are strictly conserved within orthopoxviruses.

This compact form of the holoenzyme in absence of bound template and primer strand DNA was unexpected as SAXS data suggested larger dimensions of the complex (Table 1, Fig 3AB). We hypothesized initially that we observed a compact, but inactive form of the complex induced by the sample preparation conditions, whereas the solution structure would be more open. Verification of the 2D classifications of the particles in cryo-EM gave no hint at classes with larger dimensions. In order to explore potential conformations of the complex and to assess the importance of the E9-D4 interface we first used Alphafold 2.

**Table 1.**
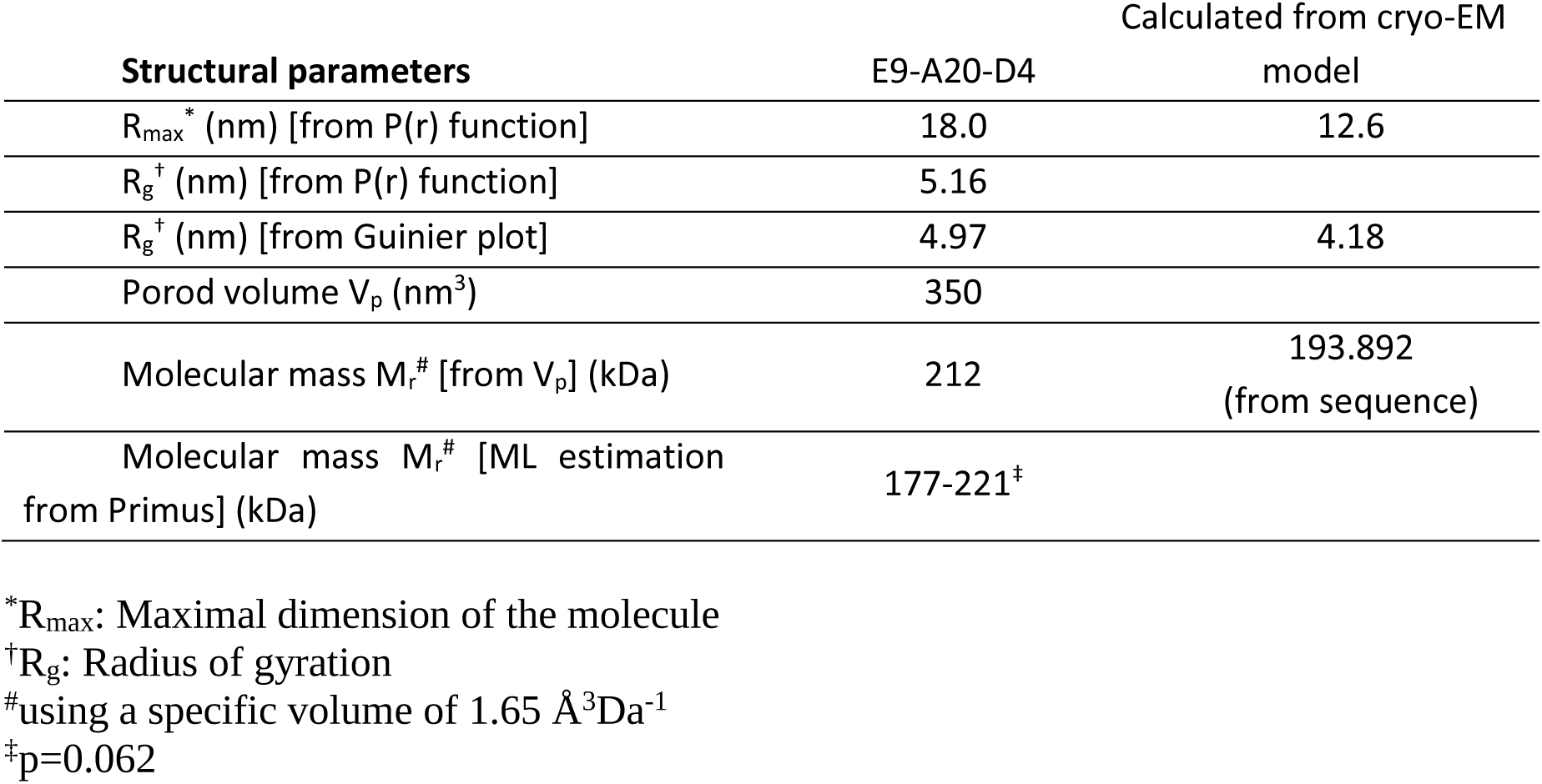
Parameters derived from SAXS measurements.

**Fig 3.**
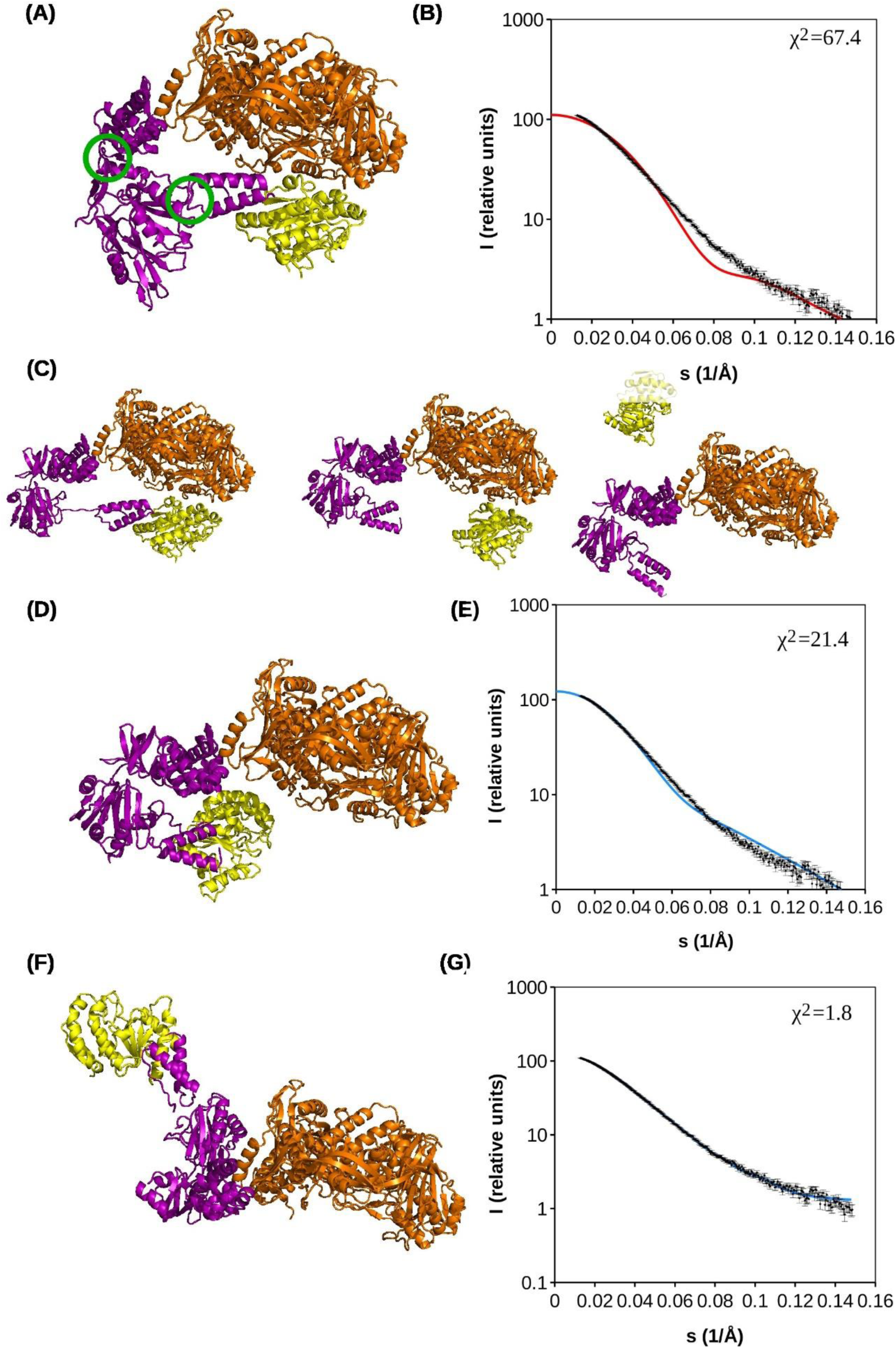
E9 holoenzyme models and their agreement with SAXS data. (**A**) Refined 3.8 Å cryo-EM structure of the holoenzyme. Green circles indicates the domain boundaries of A20, likely sites of flexible hinges in the holoenzyme. (**B**) Agreement of the experimental SAXS scattering curve and the model in (A). (**C**) Different types of solutions of multiple runs of Alphafold 2 on E9-A20-D4: models similar to the complex observed in cryo-EM (A, left), complex with an E9-D4 interaction similar to the one in (A), but with a disruption of the complex at the level of the A20-D4 interface (middle), solutions with isolated D4 (right). (**D**) The middle model of (C) is used with an added A20-D4 contact. (**E**) Calculated scattering curve of (D) compared to the experimental one. (**F**) Model of E9-A20-D4 refined with Coral against the scattering curve using the two hinges indicated in (A). (**G**) Fit of the refined model (F) against the experimental data.

When Alphafold 2 was used to calculate the structure of the holoenzyme from the sequences of the 3 involved proteins, the 25 proposed structures systematically predicted the E9-A20 interface of which the tight interaction had been demonstrated before [17], for a fraction of the structures, the A20-D4 interface has not been identified despite the tight interaction observed experimentally [15]. For a number of structures an interaction of E9 and D4 similar to the observed one has been predicted. This led to E9-A20-D4 structures, which were either similar to the cryo-EM model (Fig 3C left), to structures where the A20-D4 interaction was lost but with an interaction between E9 and D4 (Fig 3C middle) or where both interactions are lost leading to an isolated D4 subunit (Fig 3C right). Assuming that the loss of the A20-D4 interaction was not realistic, open models could be built where D4 was again associated with A20 (Fig 3D) using the known relative orientation of the two partners [15]. Such a model is already more open and explains the SAXS scattering curve better than the cryo-EM structure (Fig 3E). Using the flexibility at the two identified hinges within A20 and another hinge connecting the thumb domain to the body of E9 an almost perfect fit (χ^2^=1.82) of the scattering curve could be obtained (Fig 3FG), which agreed also in the maximal dimension and R_g_ (Table 1). This model is certainly inaccurate, as rather an ensemble of structures would contribute to the scattering curve; but with the limited amount of experimental data, we renounced from further modeling and concluded that in solution certainly more open conformations of the holoenzyme were present.

The next point to address was the importance of the E9-D4 interface. We first tried to measure a direct interaction between the polymerase subunit E9 and a monomeric mutant of D4, D4KEK. This mutant expresses better and does not form dimers in solution as wild-type D4 does [23]. Using BioLayer interferometry (BLI), it was not possible to observe an interaction between E9 and D4KEK (Fig 4A). In control experiments the previously known interaction between E9 and A20_304-426_ [16] could be confirmed (Fig 4B) as well as the interaction of E9 with a dsDNA substrate [16] (Fig 4C). We conclude that the E9-D4 interaction must be transient. In order to address the question whether the interaction is also essential, we turned to the generation of mutant VACV using a CRISPR-Cas9 based system [24] targeting the most prominent residues of the E9-D4 interface: F179 and L278 of E9 and W36 of D4 (Fig 2B) with mutations to alanine or a more radical introduction of a charged aspartic acid residue (Table 2). The approach have been validated previously on the E9-A20 interface [24], where the results were correlated with complex formation [16] so that it is expected that the number of recombinants with the expected mutation decreases with the functional impairment of the mutant. The mutation of W36 of D4 to aspartic acid (Table 2) does not allow to recover viable virus whereas the mutation to alanine seemed to be more tolerated. Also the mutation of E9 F179 to aspartic acid appeared to be fatal whereas the mutation of L278 to alanine still allowed the recovery of virus. The viable alanine mutants have a growth rate reduced by only about 20 % (S2 Fig). The effect of the mutations is in agreement with the expectations from the analysis of the 3-dimensional structure of the interface (Fig 2B) and we conclude that the E9-D4 interaction is transient, but essential.

**Table 3.**
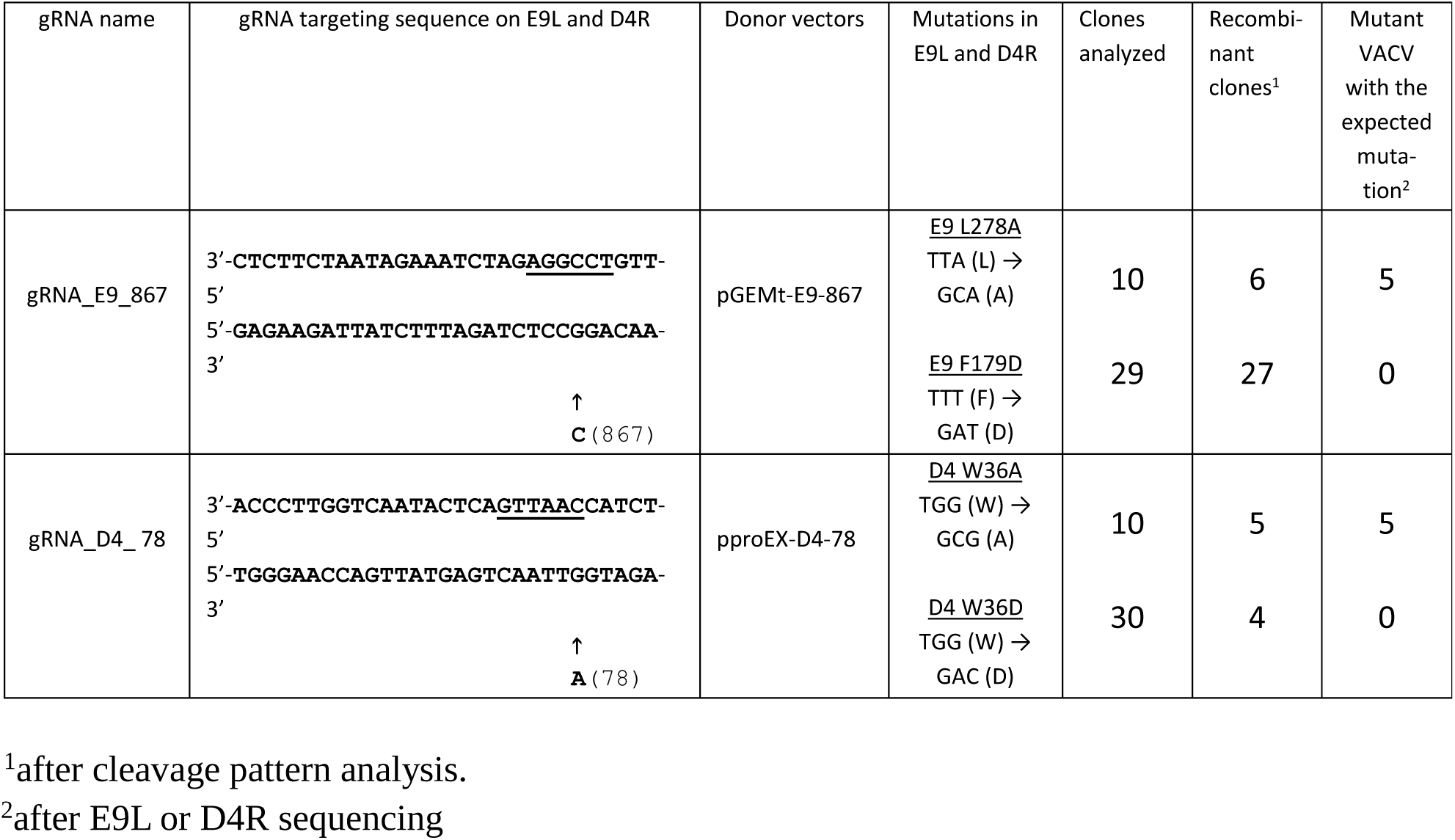
Mutant viruses generated with the CRISPR/Cas9 system.

**Fig 4.**
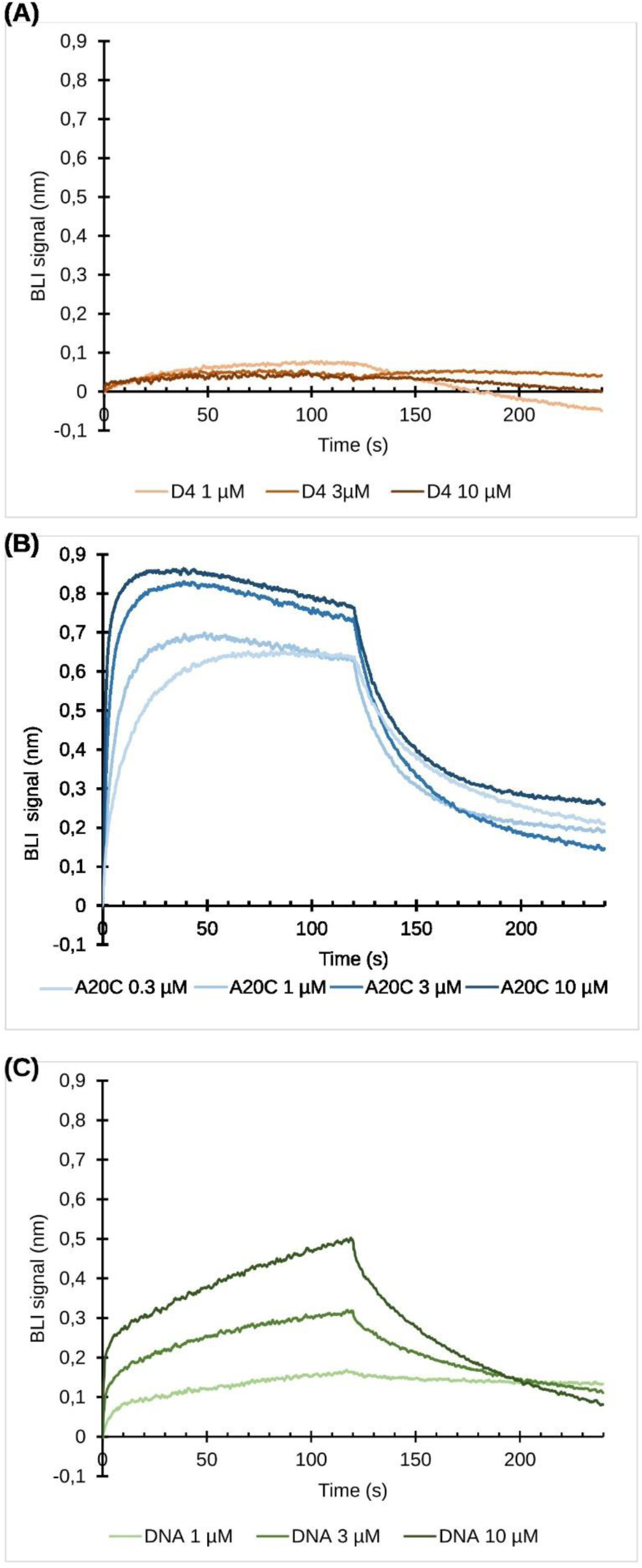
Measurements of interactions by BLI. E9 immobilized on a sensor tip using its 6His-tag is used. (**A**) Interaction with a D4 monomeric mutant (D4KEK). (**B**) Interaction with A20_304-426_. (**C**) Interaction of E9 with a dsDNA composed of a 37 base template strand with a 25 base 5’ overhang and a 12 base primer strand.

## Discussion

Following the mpox outbreak, several structures of the MPXV DNA holoenzyme became available, with [18,21] or without bound dsDNA substrate [20,21]. Surprisingly, the apo structure of the VACV polymerase holoenzyme (Fig 2A) believed initially to represent an inactive state induced by the cryo-EM sample preparation showed a closed conformation similar to the form with a bound dsDNA mimicking template strand, primer strand and incoming nucleotide [18]. In contrast to the findings by Li and co-workers [20] on MPXV holoenzyme in absence of DNA substrates, there is no indication of a formation of a dimer of heterotrimers. A superposition of the heterotrimer of VACV onto the one from MPXV [20] shows a near to identical overall structure (S3A Fig), but the ligase domain differs considerably in structure and orientation, leading to 4.58 Å rms differences for Cα positions when A20 is compared to the MPXV counterpart (S3B Fig) and still 2.28 Å rms when only the ligase domains are superposed. The electron density of Li and co-workers for this domain is far below the overall resolution of 3.1 Å and is not easily interpretable explaining why 57 residues (res.46-101) from the linker with the N-terminal domain and the N-terminal part of the ligase domain have not been modelled. As our structure of the VACV ligase domain is close to the MPXV one from Peng and coworkers and also the Alphafold model (Fig 2C), we assume that is closer to the true structure. It fits the electron density deposited in EMD-34887 evenly well as the model in pdb entry 8hm0 of Li and co-workers [20]. The residues of the E9-D4 contact are located in well-defined parts of the structures and are the same for the apo form of MPXV and VACV holoenzymes. Differences in sidechain conformations are within the error margin resulting from a limited resolution of the two structures.

A comparison of the structure of E9 and of the E9-A20-D4 complex showed the absence of any significant structural rearrangements upon complex formation, which could be analyzed due to the availability of several crystal [15,16] and NMR structures [17] of domains and proteins. The thumb domain is disordered in the cryo-EM structure whereas it is stabilized in the crystal structure due to a contact with a neighboring molecule also explaining why its orientation is very different from the commonly observed ones within family B polymerases [16]. For the body of E9, the Cα backbone in the context of the holoenzyme moves by less than 1 Å rms compared to isolated E9 (Fig 5A).

**Fig 5.**
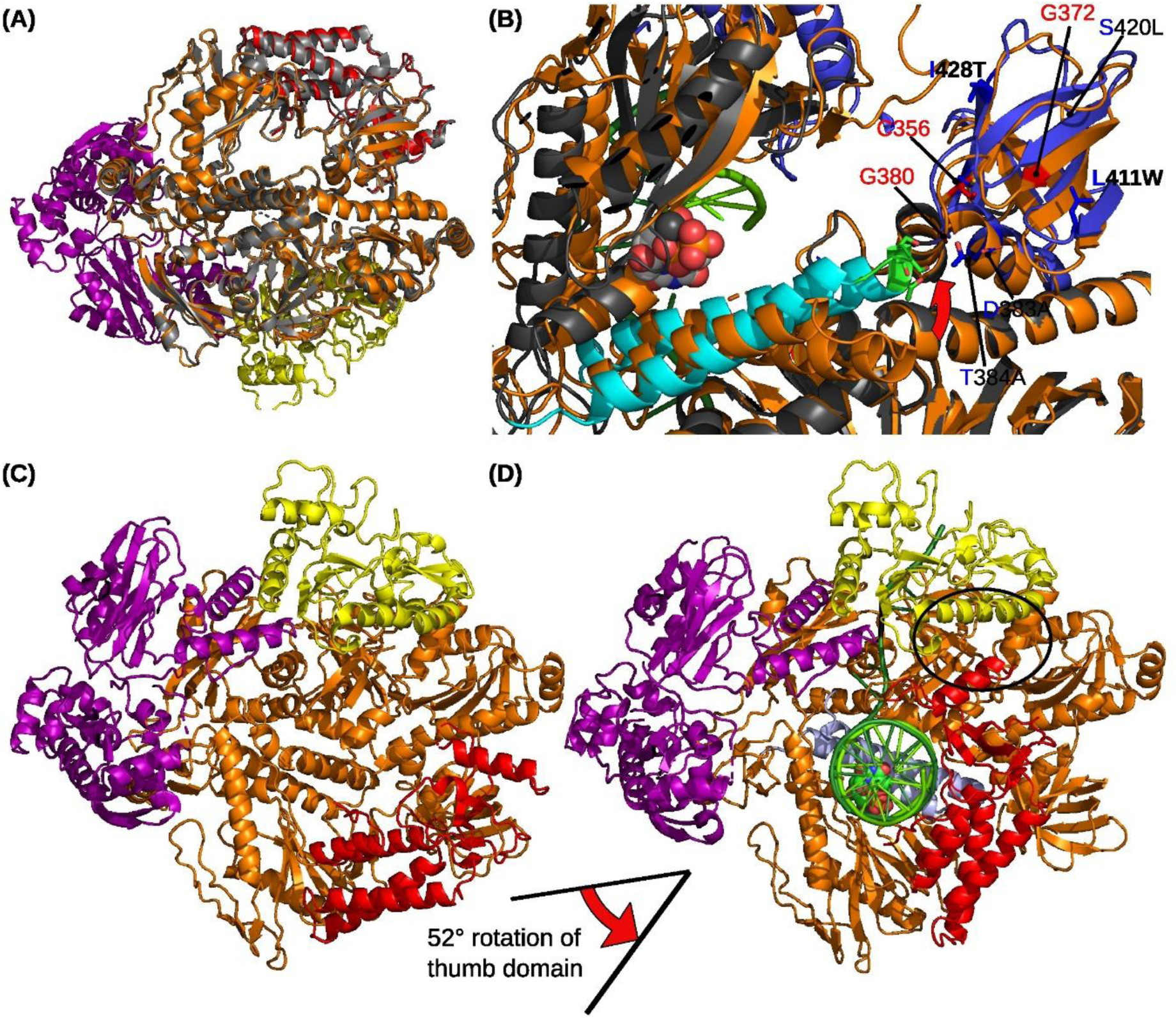
Movements in the holoenzyme complex. (**A**) E9 crystal structure (pdb 5n2e, gray) superposed onto the refined structure of the E9-A20-D4 complex (colors as in Fig 2). (**B**) Close-up of the fingertip and the insert 2 region of the apo form of VACV holoenzyme (orange) superposed onto the DNA-bound MPXV F8 ([19], gray, finger in cyan, residues of the fingertip in stick representation and highlighted in green, insert 2 domain in blue) with an incoming dCTP nucleotide (spheres). A red arrow shows the direction of the movement. Residues involved in PAA resistance of VACV polymerase [49] are labeled and shown in red. Residues of insert 2 differing between MPXV (GenBank accession ON755039, [48]) and VACV are shown in stick representation and labelled (the residues of F8 are printed in blue, the ones of E9 in black) and recently acquired mutations are printed in bold. (**C**) VACV E9-A20-D4 with the disordered thumb domain in red positioned according to the crystal structure. (**D**) Same orientation of the MPXV holoenzyme with bound DNA and incoming dTTP nucleotide. Domains are colored as in (C), with the DNA in green and dTTP shown as spheres. The contact area created between the thumb domain of F8 and E4 is indicated by an ellipse.

The 3-dimensionnal structure of the VACV holoenzyme allows to revisit previous work [25] that had identified several temperature-sensitive (ts) mutants of A20 compromising processive DNA replication while conserving the distributive action of E9 (S3 Table). Punjabi and coworkers [25] analyzed previously identified ts mutants and created a number of clustered charged-to-alanine mutants. Similar charged-to-alanine VACV mutants have been analyzed for viral growth by Ishii and co-workers [26] (S3 Table). The single mutant Dts48 with a G84E mutation [27] is situated in the connection between N-terminal and ligase domain, the mutation A20-ER-5 located at the level of res. 185-191 of the ligase subdomain and A20-6 affecting residues 265-269 of the OB domain at the interface with the ligase subdomain [9,25]. Whereas the first mutation might affect the flexibility of A20, the ts-mutants would probably rather lead to misfolding of A20 at the non-permissive temperature and interfere with the assembly of a functional trimer. Two of the charged-to-alanine mutants of Punjabi and coworkers [25] were inactive in processive DNA synthesis, again probably through misfolding, explained by a role of D178 of the 177 DDE to AAA mutant in the capping of a small helix and of E185 of the 185ER to AA mutant in structural hydrogen bonds. Why the more radical A20-ER-5 mutant only led to the above mentioned ts phenotype remains obscure but agrees with the results of Ishii et *al*. [26]. Globally, there is a very good correlation between the effect of the mutation in processive DNA synthesis and viral growth with the structural role of the residues (S3 Table), although the effect of mutants introducing multiple alanine residues at the same time may affect non-specifically the secondary structure leading to inactive or ts mutants.

The dimer of trimers observed by Li and coworkers [20] in pdb entry 8hlz is intriguing. The assembly of this MPXV hexamer if formed by two extended F8-A22-E4 complexes interacting around a quasi-2-fold axis using in *trans* the E9-D4 interface observed in the heterotrimer (S3C Fig, top) creating additional contacts between thumb domain and A22. In order to explain the transition to a compact trimer with an E9-D4 contact in *cis*, the author proposed a dissociation of A22 from D4 in *trans* position and rebinding of A22 to D4 in cis. This model can be refuted as we showed that the E9-D4 interaction is transient with less important buried surface (380 Å^2^ analyzed with PDBePISA [28]) than the A20-D4 interaction (830 Å^2^) or for comparison the E9-A20 interface (670 Å^2^). In addition, the interface appears to be extremely important as it is not possible to produce A20 without bound D4 [15]. We rather propose a transition of the hexamer to the trimer involving a dissociation of E4 from F8 in the hexamer, a change of the conformation of A22 and rebinding to F8 in *cis* leading to two compact trimeric complexes (pdb entry 8hm0, S3C Fig, bottom).

The DNA-free cryo-EM structure of VACV E9-A20-D4 clearly exhibited discrepancies when compared to the SAXS data from the holoenzyme complex in solution (as shown in Fig 3B). This suggested a distinct conformation with a greater radius of gyration (4.97 nm vs. 4.18 nm) and a larger maximal dimension (18.0 nm vs. 12.6 nm, as outlined in Table 1).

For A20, Alphafold 2 predicted 3 domains and allowed to identify two hinge regions (Fig 2C) introducing flexibility, the first one in the linker of res. 50 to 67 separating the N-terminal domain from the ligase domain and the second hinge around residue 310, in the connection between ligase domain and the C-terminal domain A20_304-426_. This flexibility allowed to fit the Alphafold 2 model of A20 into the cryo-EM electron density without further modifications (Fig 2AC). Introducing flexible linkers into the model allowed to fit the SAXS curve unexpectedly well (Fig 3FG), although in reality rather an ensemble of different conformations including also closed ones, is expected.

The existence of extended and closed conformations is also supported by the explanation of the SAXS scattering curve by a combination of an extended VACV holoenzyme model (S3C Fig) based on the MPXV hexamer structure [20] and the compact holoenzyme of our structure (S4 Fig). The maximal dimension of 18 nm of the extended model (S4A Fig) corresponds to the maximal dimension of the VACV virus holoenzyme complex in solution determined by SAXS (Table 2). The mixture of compact (42 %) and the extended (58 %) VACV holoenzyme fitted the observed SAXS scattering curve (S4C Fig) as well as the Coral-fitted model (Fig 3FG).

The conformational variability of the holoenzyme has been explored further by Alphafold 2 predictions of the complex structure. The solutions were either similar to the closed form (Fig 3C, left) or to open forms (Fig 3C, right). We concluded that open conformations are present in solution, but had still to verify the importance of the closed conformation and the strength of the E9-D4 interface. Previous publications had identified the E9-A20 interaction and the A20-D4 interaction but never an E9-D4 interaction [9] and indeed the interface area (380 Å^2^) of the E9-D4 interface is much smaller than the one of A20-D4 (830 Å^2^) and E9-A20 (670 Å^2^). It was not possible to measure an affinity of the direct interaction of E9 and D4 in contrast to the established interactions of E9 with A20_304-426_ and with DNA underlining the transient character of the interaction (Fig 4). Still, site-directed mutagenesis results (Table 2) showed that viruses mutated radically in key residues of the interface are not viable highlighting the importance of the interaction, whereas less drastic mutations seem to be tolerated.

The comparison with the DNA bound structure of Peng and co-workers [18] shows a very similar conformation of the holoenzyme and allowed to understand the role of the E9-D4 interaction (Fig 5CD) leading to the formation of a ring entrapping the template strand. The neo-synthesized primer strand leads to a movement of the thumb domain, which closes upon the dsDNA leading to a rotation of 52° relative to the orientation in the E9 crystal structure (Fig 5C). This movement brings residues 896-925 inserted into the 4-helix bundle of the thumb domain next to D4, assigning indirectly a role for this extension conserved within orthopoxviruses. It appears to form a platform supporting D4 but without direct E9-D4 interactions, which are restricted to the previously analyzed hydrophobic interface around tryptophan 34 of D4 (Fig 2B). The thumb domain closes onto the neo-synthesized dsDNA whereas the E9 - D4 contact encircles the template DNA assuring the processivity of the polymerase.

At last, a structural difference between apo and DNA bound holoenzyme is the movement of the fingertip observed in the MPXV structure in presence of DNA and an incoming dTTP nucleotide, which is characteristic for class B polymerases [29,30] and has been proposed in the quest for a role of the insert 2 domain (res. 356–432) [16]. Indeed, the tip of the finger domain contacts insert 2 in the MPXV structure as predicted for VACV [16] (Fig 5B). The role of this interaction is not understood but insert 2 carries resistance mutations against phosphonoacetic acid (PAA), a pyrophosphate analog inhibiting VACV polymerase (shown in red in Fig 5B). Insert 2 also clusters sequence differences between VACV Copenhagen and MPXV virus (Zaire-96-I-16, NCBI NP_536484.1, Fig 5B, black labels) including some recently acquired mutations in isolate IRBA22-11 [31] (Fig 5B, bold black labels).

The model presented by Peng and co-workers [18] is very suggestive regarding the position of D4 as the template strand appears to continue into the DNA binding site of D4 as identified by a crystal structure with bound dsDNA comprising an abasic site [19], where we had also shown that D4 can excise uracil bases in the context of single-stranded or double stranded DNA. This speaks in favor of an action of UNG on the template strand and stalling of the polymerase upon the detection of a uracil residue in the template strand has indeed been observed [27]. But a close inspection of the published electron density [18] shows that there is a break in the backbone density between the DNA bound to the polymerase and the DNA bound to the UNG active site where the density is rather disordered raising doubts about the continuity of the observed DNA. As in presence of dUTP, abasic sites occur in the neo-synthesized strand [27], this is an argument in favor of a different positioning of D4 downstream of the polymerase active site. Future structural and biochemical work is needed in order to validate one of these hypotheses and also to cast light on the role of E9 as recombinase, which uses only the exonuclease site and neither the polymerase active site nor the processivity factors.

## Materials and Methods

### Construction of the baculovirus expressing E9-A20-D4

The MultiBac baculovirus/insect cell expression system [32] was used in order to express the VACV DNA polymerase holoenzyme using sequences of *E9L, A20R*, and *D4R* of VACV strain Copenhagen (GenBank accession number <u>M35027.1</u>). Gene synthesis and cloning was performed by GenScript. *E9L* was inserted into the pUCDM vector. Recombinant E9 contains a 6His-tag and the Tobacco Etch Virus (TEV) protease cleavage site sequence fused to its N-terminus. *A20R* and *D4R* (D4 carrying an N-terminal Strep-tag) were both cloned into the pFL plasmid. In consequence, E9 carries an N-terminal TEV-cleavable His-Tag with the sequence MSYYHHHHHH DYDIPTTENL YFQ↓GAMDP, which replaces the N-terminal methionine. D4 carries an N-terminal Strep-tag with the sequence MASWSHPQFE KSGGGGGLVP RGSA before the N-terminal methionine residue.

*E9L, A20R*, and *D4R* were then combined by *in vitro* fusion of acceptor (pFL) and donor (pUCDM) plasmid derivatives using Cre recombinase [33]. The resulting construct was used to transform DH10EMBacY bacteria provided by the Eukaryotic Expression Facility (EEF, EMBL Grenoble). Bacmid carrying the VACV genes was extracted, purified and transfected into *S. frugiperda* (*Sf21*) cells. Transfection of 10^6^ cells in a 6-well plate was performed using 5 μl of transfection agent (X-tremeGENE HP Reagent, Roche), 10 μL of bacmid and 85 μL of SF900II-SFM medium (Gibco). *Sf21* cells were incubated for 48 h at 27 °C. The virus-containing supernatant (V_0_) was recovered. The viral stock for protein expression was prepared as follows: 25 mL of *Sf21* cells at 0.5×10^6^ cells/mL were infected with 3 mL of V_0_. Cells were maintained at 0.5·106 cells/mL until cell growth stopped. After centrifugation at 1000×g for 3 min, the supernatant containing recombinant baculovirus was recovered. Viral stocks were stored at 4 °C, protected from light.

### Protein expression and production

E9-A20-D4 has been purified using a long protocol or a simplified protocol. Frozen cell pellets from 2 L of infected High Five (Thermo Scientific) insect cell culture were resuspended in 50 mM Tris-HCl pH 8.5, 300 mM NaCl, 10 % glycerol, 5 mM imidazole, 5 mM β-mercaptoethanol (buffer A) with Complete protease inhibitor cocktail (Roche) and 2 mg DNAse I from bovine pancreas (Sigma-Aldrich) and lysed in a Potter homogenizer (B. Braun, Melsungen, Germany) using 20 strokes or by sonication (5 min, Cycle: 0.5, Amplitude: 70 %; Labsonic®, Satorius). After centrifugation (58 000×g at 4 °C) for 20 min, the supernatant was loaded on a 5 mL His-trap FF Crude column (Cytiva) using a peristaltic pump. The column was washed with 20 mL of buffer A and eluted with 20 mL of the same buffer containing 180 mM imidazole.

The sample was concentrated on a Vivaspin 30 kDa centrifugal concentrator (Sartorius) to 3 mL for buffer exchange with an Econo10 DG column (Biorad) equilibrated in buffer A and eluted with 4 mL of the same buffer. 1/100 (w/w) TEV protease were added and incubated overnight at room temperature.

The digested complex was diluted to 15 mL with buffer A and loaded on same Ni-column equilibrated in buffer A. The column was washed with 15 mL of buffer A first, followed by 20 mL of 20 mM imidazole in buffer A. Complex-containing fractions were pooled, concentrated to 3.5 mL and diluted with 50 mM Tris-Hcl pH 8.5 for a final NaCl concentration of 200 mM. A 1.5 mL Streptactin (IBA Lifesciences, Göttingen, Germany) column was equilibrated in 100 mM Tris-HCl pH 8.5, 200 mM NaCl (buffer C), the sample was loaded and the column washed with 5 mL buffer C. The complex was eluted in five 1 mL fractions using buffer C with added 5 mM desthiobiotin (IBA Lifesciences). After addition of TCEP to 1 mM, complex containing fractions were concentrated to 0.8 mL and injected on a Superdex S200 10/300 (Cytiva) column equilibrated in 20 mM Tris-HCl pH 8.5, 200 mM NaCl and 1 mM TCEP. The peak fractions were concentrated to ≈ 2 mg·mL^-1^ for further use in structural studies.

A simplified protocol has been used for the cryo-EM studies. It shunts the TEV digestion and the second Ni-column and used the buffer-exchanged protein directly on the streptactin column omitting also the size exclusion chromatography step.

### Multiple Angle Light Scattering (MALS)

SEC was performed with a Superdex 200 10/300 GL (GE Healthcare) equilibrated in 50 mM Tris-HCl pH 7.5, 100 mM NaCl. The run was performed at 20 °C with a flow rate of 0.5 mL·min^−1^. 50 μL of a protein solution at a concentration of 2 mg·mL^−1^ were injected. On-line MALS detection was performed with a DAWN-EOS detector (Wyatt Technology Corp., Santa Barbara, CA) using a laser emitting at 690 nm. The protein concentration was measured on-line by refractive index measurements using a RI2000 detector (Schambeck SFD) using a refractive index increment dn/dc = 0.185 mL·g^-1^. Data were analyzed and weight-averaged molecular masses (Mw) were calculated using the software ASTRA V (Wyatt Technology Corp., Santa Barbara, CA) as described [34].

#### SEC-SAXS

SAXS measurements were done on BM29 of ESRF at a ring current of 200 mA. 45 μL of E9-A20-D4 complex with a cleaved His-tag at 2 mg·mL^-1^ in 25 mM Tris-HCl pH 7.5, 300 mM NaCl were injected onto a Superdex S200 3.2/100 (Cytiva) size exclusion column. Runs were performed at a flow rate of 0.1 mL·min^-1^ and 2000 frames of 1 s were collected using a Pilatus 1M detector (Dectris). Individual frames were processed automatically and independently within the EDNA framework yielding radially averaged curves of normalized intensity versus scattering angle s=4πsinθ/λ [35]. Frames corresponding to the elution of E9-A20-D4 were identified in iSPyB [36], merged and analyzed further using Primus of the ATSAS package [37].

#### SAXS data treatment

SAXS data were analyzed with different softwares of the ATSAS package [37]. Scattering curves of models were calculated and compared to the experimental scattering curve using Crysol. Rigid-body modeling with Coral used 4 bodies (D4+A20_1-54_, A20 ligase domain (res. 56-310), A20_312-426_+E9_1-830_, E9 thumb domain E9_832-1006_) based on the E9-A20-D4 model from cryo-EM. A linker residue has been introduced at the position of the hinges of A20 and the connection of the thumb to the body of the polymerase. The relative contributions of extended and compact forms of the holoenzyme were obtained with Oligomer [38].

#### Alphafold 2 predictions

Predictions of the complex structure used an Alphafold 2 [39] installation on the CCRT-HPC TOPAZE super calculator from the CEA (https://www-ccrt.cea.fr). The sequences of the three proteins of E9-A20-D4 have been provided and 5 runs of 5 models have been generated resulting in an ensemble of 25 predicted conformations of the trimeric complex.

#### Cryo-EM of the apo form of the E9-A20-D4 holoenzyme

The purification omitted TEV cleavage and the 2^nd^ Ni-column purification step and E9-A20-D4 complex was used directly after the Streptactin column purification step and concentrated in a Vivaspin 6 concentrator with 30 kDa cut-off to 1.7 mg·mL^-1^. The sample was centrifuged for 20 min at 13 000 rpm in an Eppendorf centrifuge. 5 μL of a buffer (80 mM Tris-HCl pH 7.4, 20 mM EDTA) were added to 15 μL of complex. In the Vitrobot chamber with 100 % humidity at 4 °C, 4 μL of sample were applied for 30 s to 1.2/1.3 holey carbon on 300 mesh copper grids (Quantifoil), followed by 5 s blotting with blot force 20 and plunge freezing in liquid ethane with a Vitrobot IV (FEI, Thermo Scientific). Grids were glow discharged for 90 s in air at 13 Pa in a Plasma Cleaner model PDC-002 (Harrick Plasma, Ithaca, NY, USA).

#### Cryo-EM data collection and processing

Data were collected with EPU on the FEI Krios electron microscope (Thermo Scientific) of the Rudolf-Virchow-Zentrum, Würzburg, Germany. Statistics are given in Table 1. Data were processed on the EMBL-IBS computing cluster with a SBGrid [40] software installation. Motion correction used Relion 4.0 [41]. The downstream data processing used Cryosparc [42] using the pipeline shown in S1 Fig. The final step of non-uniform refinement refined also tilt and per group CTF parameters. Maps and models were displayed with Chimera [43], which was also used for the docking of the existing structures into the cryo-EM map: (A20_1-50_-D4 complex, pdb entry 4od8, E9-A20_304-426,_ PDB-DEV database accession code: PDBDEV_00000075, MPXV ligase domain from pdb entry 8hg1 [18]). Coot [44] was used to model the sequence of VACV A20 onto the MPXV A22 structure and for the adjustment of the linkers between individual domains. The structure was refined using Phenix real-space refinement [45] at a resolution of 3.8 Å against a map sharpened with a temperature factor of −160 Å^2^. Statistics of the final model are given in S1 Table.

#### Biolayer interferometry (BLI) measurements

BLI measurements were performed on a BLItz instrument (Forté Bio) with NTA biosensors (Sartorius) using the following protocol after at least 20 min initial rehydration: baseline measurement 60 s, E9 loading 180 s, D4 or A20 or DNA association 180 s, dissociation 180 s, regeneration 30 s, baseline 60 s, nickel loading 60 s, baseline 30 s. Running buffer was 20 mM HEPES pH 7.5, 100 mM NaCl, 0.05 % TWEEN 20. E9 with an uncleaved His-tag [16] was used at 0.1 mg·mL^-1^ for loading; analyte concentrations varied from 0.03 to 10 μM, regeneration used 10 mM glycine pH 1.7 buffer and nickel loading used 10 mM NiCl_2_. Data were baseline corrected, normalized for the quantity of E9 bound to the chip and corrected for dissociation of His-bound E9 using Excel (microsoft.com).

#### D4-KEK mutant

In order to obtain a D4 construct, which does not dimerize at high concentration as described by Schormann and co-workers [23], we modified residues in the D4-D4 interface. Hydrophobic residues I197, V200 and L204 involved in the D4-D4 and D4-A20 interfaces were changed to their charged counterparts in the human UNG protein, which was known to be strictly monomeric. The “humanized” mutant D4 I197K-V200E-L204K (D4KEK) was obtained using the previously described pPROEX-D4 plasmid [15] mutagenized by PCR amplification of the full plasmid using primers ttCGTTGATt tTTTCAAATG ATCTATCTTT CTCG and TTACTGGAAa aAGACAACAA GGTACCTATA AATTGG (lower letters indicate the introduced mutations) followed by recircularization by ligation. The resulting construct contains an N-terminal 6His-tag and a TEV cleavage site. D4KEK was expressed and purified using the same protocol as described for wild-type D4 [15]. Initial crystallization conditions of D4KEK were obtained using the EMBL HTX platform and were refined manually to 22 % PEG4000, 100 mM LiSO_4_, 100 mM Hepes pH 7.5. Diffraction data were collected on the SOLEIL PX1 beamline on 25/07/2019. Data were reduced with XDS [46], analyzed with AIMLESS [47] and molecular replacement solution was found with MOLREP [48] using the wild type D4 structure (chain B of pdb 4od8). The structure was refined to 1.43Å resolution using cycles of refinement with REFMAC [49] and manual building with COOT [44]. The final model, superposes very well on wt D4 (0.25 Å rmsd on Cα atoms) and statistics are given in S2 Table.

#### Production of recombinant VACV carrying mutations in E9L and D4R

The production of mutant VACV using the CRISPR/Cas9 mediated homologous recombination was described in Boutin *et al*. [24]. Briefly, 5×10^6^ CV-1 cells were infected with VACV at a MOI of 0.02 in DMEM supplemented with 0.5 % *v/v* FCS for 1 h at 37 °C in 5 % CO_2_ atmosphere. Cells were then electroporated using the Neon Transfection System (Thermofisher) as follows: cells were washed once in 1× phosphate-buffered saline (PBS), harvested by trypsinization, and resuspended in 100 μL of the kits electroporation buffer R (1×10^7^ cells/mL). Cells (100 μL) were mixed with 2.25 μg of each plasmid: pCMV-Cas9ΔNLS, a donor vector carrying the mutated *E9L* or *D4R* and pU6-gRNA (Sigma) encoding gRNA targeting *E9L* or *D4R* (Table 3). The cell/DNA mixture was aspirated into a 100 μL Neon tip and submitted to two electric pulses at 1050 V for 30 ms. Cells were then seeded in six-well plates containing warm DMEM supplemented with 10 % FCS and incubated for three days at 37 °C in 5 % CO_2_. After a single freeze-thaw cycle, the viral suspension was recovered and diluted in DMEM before infection of Vero cells seeded in six-well plates. After 1 h of adsorption, the residual inoculum was removed and replaced with DMEM supplemented with 10 % FCS and 0.6 % agarose. The plates were incubated at 37 °C in 5 % CO_2_. Three days post-infection, 10 to 20 individual plaques were picked and used to infect Vero cells seeded in 96-well plates for two days. For each clone, the viral DNA genome was extracted using the QIAamp DNA mini kit (Qiagen). Specific genomic regions of *E9L* or *D4R* were PCR-amplified using the following primers: E9-F: 5’CTAACAAAGA GCGACGTACA AC3’, E9-R: 5’GAAGCCGTCG ATAGAGGATG3’ and D4-F: 5’CTATAGGACC TTCCAACTG3’, D4-R: 5’CCTTGAGCCC AATTTATAGG3’, respectively. PCR amplicons were purified using a QIAquick purification PCR kit (Qiagen) and digested with BspEI (*E9L*) or MfeI (*D4R*) restriction enzymes (NEB). Digested products were separated by 0.8 % agarose gel electrophoresis and visualized with ethidium bromide. Mutant viruses were further characterized by Sanger sequencing of the *E9L* or *D4R* gene to ensure the presence of the expected mutations.

**Table 3.** Cryo-EM data collection statistics.

| **Parameter** | **Cryo-EM data collection**<br>**E9-A20-D4** |
| --- | --- |
| Microscope | Titan Krios G3 (Thermo Fisher) |
| High tension (kV) | 300 |
| C<sub>s</sub> (mm) | 2,7 |
| Detector | Falcon III camera |
| Mode | Counting |
| Energy filter | No |
| Magnification | 42000 x |
| Data collection date | 28/2/2022 |
| Calibrated pixel size | 1.064 Å |
| No. of frames per movie | 47 |
| Nominal defocus range (µm) | - (1.4–2.4) |
| Dose par frame (e·Å<sup>-2</sup>) | 1.45 |
| Dose Rate (e·px<sup>-1</sup>·s<sup>-1</sup>) | 1.03 |
| Drift rate (nm·s<sup>-1</sup>) | < 0.04 |
| No. of movies | 1376 |

#### Growth kinetics of mutant VACV in vitro

Twenty-four well plates containing 8×10^5^ Vero cells were infected with VACV at a MOI of 0.05 in DMEM at 37 °C in a 5 % CO_2_ atmosphere. At 1 h post infection (hpi), the viral inoculum is removed and fresh media containing 2.5 % v/v FCS was added and the plates incubated at 37 °C. Cells were harvested at 8, 24, 48, and 72 hpi. Virus yield was determined by titration of the virus on Vero cells as described [50].

## Acknowledgments

We thank Guy Schoehn for data collections on the Glacios electron microscope and his operation of the IBS EM facility. We are grateful to Aymeric Puech and Rémi Pinck for assistance and operation of the IBS-EMBL cryo-EM computing facility. WPB acknowledges the support of the CNRS for a sabbatical in Utz Fischer’s group at the Biozentrum, University of Würzburg. WPB is grateful to Emilia Gärtner for excellent support. We thank Fred Garzoni and Imre Berger for help in setting up the Multibac system.

## Data availability

The cryo-EM density map has been deposited in EMDB entry EMD-18134, the corresponding model in PDB entry 8q3r. The structure of the D4 mutant D4KEK has be deposited in pdb entry 8qam. SAXS data have been deposited in SASBDB entry SASDSW8.

## Supporting information

**S1 Table. Cryo-EM map and model statistics**

**S2 Table. Data collection and model statistics for the D5KEK crystal structure**

**S3 Table. Structure – activity relationship of previously described charged-to-alanine mutants in A20**

**S1 Fig. Cryo-EM structure determination**.

**S2 Fig. Growth kinetics of VACV mutated in the E9-D4 interface**.

**S3 Fig. Comparison of the holoenzyme heterotrimer of VACV polymerase with the one of MPXV**.

**S4 Fig. Explanation of the SAXS curve of VACV holoenzyme**.

