## Supporting information for "Structure and flexibility of the DNA polymerase holoenzyme of vaccinia virus"

**for**

Short title: VACV DNA polymerase holoenzyme

**S1 Table. Cryo-EM map and model statistics**

| **Structural parameters** | E9-A20-D4 holoenzyme without thumb domain |
|---|---|
| Resolution for FSC = 0.143 (Å) | 3.8 |
| Map Wilson factor ($Å^2$) | 180 |
| Number of non-H atoms | 11933 |
| ***MolProbity*** | |
| Model map correlation (masked) | 0.54 |
| MolProbity score | 2.9 |
| Clash score | 19.8 |
| Rms deviation from ideality, bonds (Å) | 0.004 |
| Rms deviation from ideality, angles (°) | 0.59 |
| Ramachandran plot outliers (%) | 0.3 |
| Ramachandran plot allowed (%) | 6.3 |
| Ramachandran plot favored (%) | 93.5 |

**S2 Table. Data collection and model statistics for the D5KEK crystal structure**

| | D4KEK overall | D4KEK outer shell |
|---|---|---|
| **Data collection :** | | |
| Instrument | SOLEIL PX1 | |
| Wavelength (Å) | 0.97856 | |
| Space group | I4 | |
| Cell dimensions<br>a, b, c (Å)<br>α, β, γ (°) | <br>161.67 161.67 39.28<br>90 90 90 | |
| Resolution range | 40.42-1.32 | 1.34-1.32 |
| $R_{merge}$ | 0.05 | 1.134 |
| I/σI | 23.4 | 1.9 |
| Completeness (%) | 99.5 | 90.2 |
| Total reflection | 1589908 | 63507 |
| Unique reflection | 119652 | 5311 |
| CC(1/2) | 0.999 | 0.844 |
| **Refinement** | | |
| $R_{work}/R_{free}$ | 0.129 / 0.165 | |
| No non-H atoms | 4651 | |
| No water | 738 | |
| RMSD bond length (Å) | 0.017 | |
| RMSD bond angle (°) | 2.118 | |
| Ramachandran plot outlier (%) | 0.55 | |
| Ramachandran plot allowed (%) | 2.20 | |
| Ramachandran plot favored (%) | 97.25 | |

**S3 Table. Structure – activity relationship of previously described charged-to-alanine mutants in A20**

| | **Ishii *et al.*, 2001** | | | |
|---|---|---|---|---|
| **Name** | **First residue** | **Targeted res.** | **Activity/growth** | **Explanation** |
| | 62 | DEVKNK* | no effect | Linker between N-terminal and ligase domains |
| | 108 | DDMR | no effect | Surface of ligase subdomain |
| | 167 | EIEIEED | inactive | Not obvious, affecting probably secondary structure |
| | 177 | DDE | weak ts | Helix capping by D178 |
| | 185 | ERSFDDK | stringent ts | Structural hydrogen bonds of E185 |
| | 204 | ELRR | inactive | Structural role of R207 |
| | 224 | KVDR | inactive | Not obvious, affecting probably secondary structure |
| | 248 | KDVDH | stringent ts | Structural role of H353 |
| | 255 | RSKVREH | inactive | Not obvious, K257 and E260 may have structural role |
| | 265 | KVKKK | weak ts | K269 has structural role in the OB domain - ligase subdomain interface |
| | 345 | KRKIK | weak ts | Surface of C-terminal domain, affecting potentially secondary structure |
| | **Punjabi *et al.*, 2001** | | | |
| Dts48 | 84 | Single point mutation G84E | stringent ts | Connection between N-terminal and ligase domain affecting potentially flexibility |
| 1 | 62 | DEVK | no effect | Linker between N-terminal and ligase domains |
| 2 | 108 | DDMR | no effect | Surface of ligase subdomain |
| 3 | 171 | EED | no effect | Surface residues in loop |
| 4 | 177 | DDE | inactive | Helix capping by D178 |
| 5 | 189 | DDK | no effect | Surface residues in loop |
| 6 | 265 | KVKKK | stringent ts | K269 has structural role in the OB domain - ligase subdomain interface |
| 7 | 345 | KRK | no effect | Surface of C-terminal domain |
| 8 | 345 | KRKIK | no effect | Surface of C-terminal domain |
| ER | 185 | ER | inactive | Structural hydrogen bonds of E185 in the ligase subdomain |
| ER-5 | 185 | ERSFDDK | stringent ts | Structural hydrogen bonds of E185 in ligase subdomain |

*Mutated residues are shown in color according to their charge

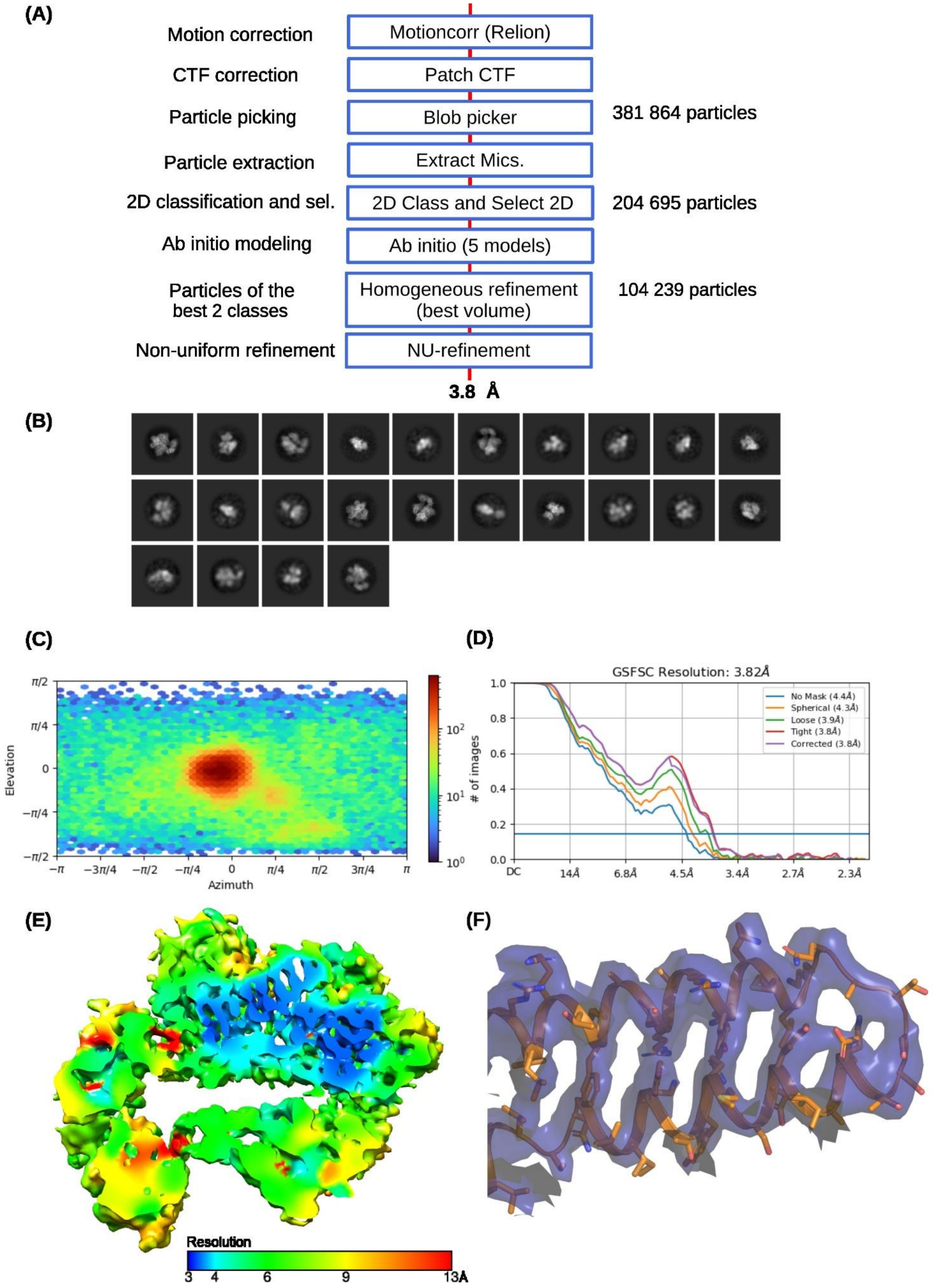

**S1 Fig. Cryo-EM structure determination.** (**A**) Flow-diagram of the structure determination using CryoSparc. (**B**) 2D classes containing 204 695 particles used for the 3D reconstruction. (**C**) Orientation of the particles used in the final model obtained after non-uniform refinement. (**D**) Gold standard Fourier shell correlation of the refined model. (**E**) Cut-away of the sharpened electron density map colored according to the local resolution. (**F**) Example electron density of the finger domain of E9 with the underlying structure in stick representation.

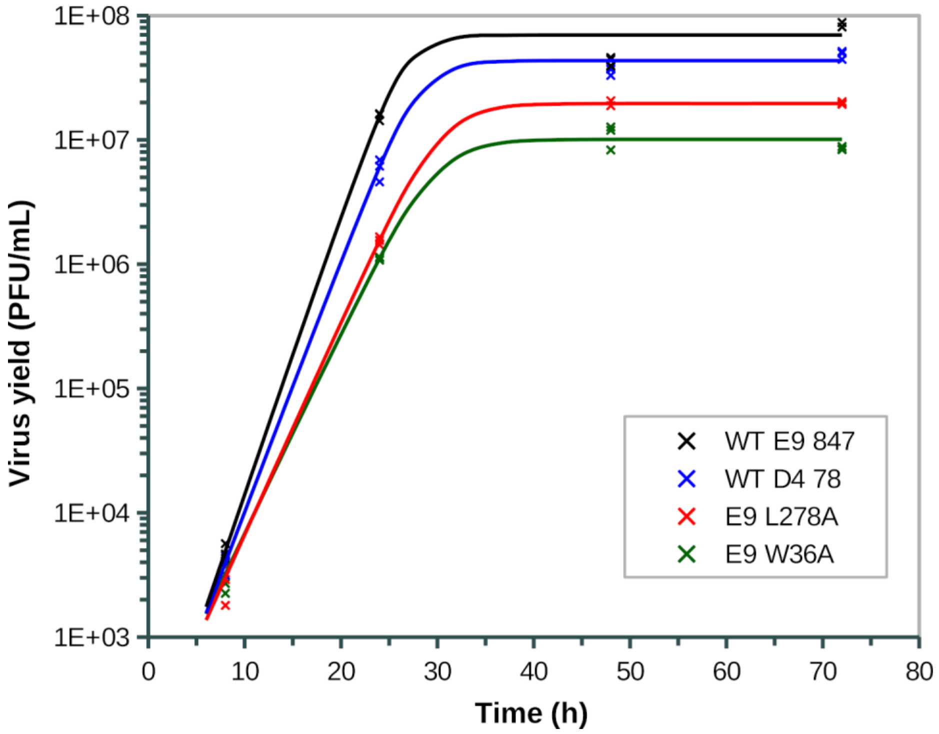

**S2 Fig. Growth kinetics of VACV mutated in the E9-D4 interface**. Vero cells were infected at a MOI of 0.05 and harvested at 8, 24, 48, and 72 hours post-infection. Viral titers were determined by plaque assays and expressed in PFU/mL. Data is representative of three independent experiments. Titrations are done in duplicate and the average is plotted. WT E9 867 and WT D4 78 are wt VACV carrying a silent mutation in E9 and D4, respectively. E9 L278A and D4 W36A are mutant VACV with mutation at the E9-D4 interface, respectively. The data points have been fitted with logistic functions.

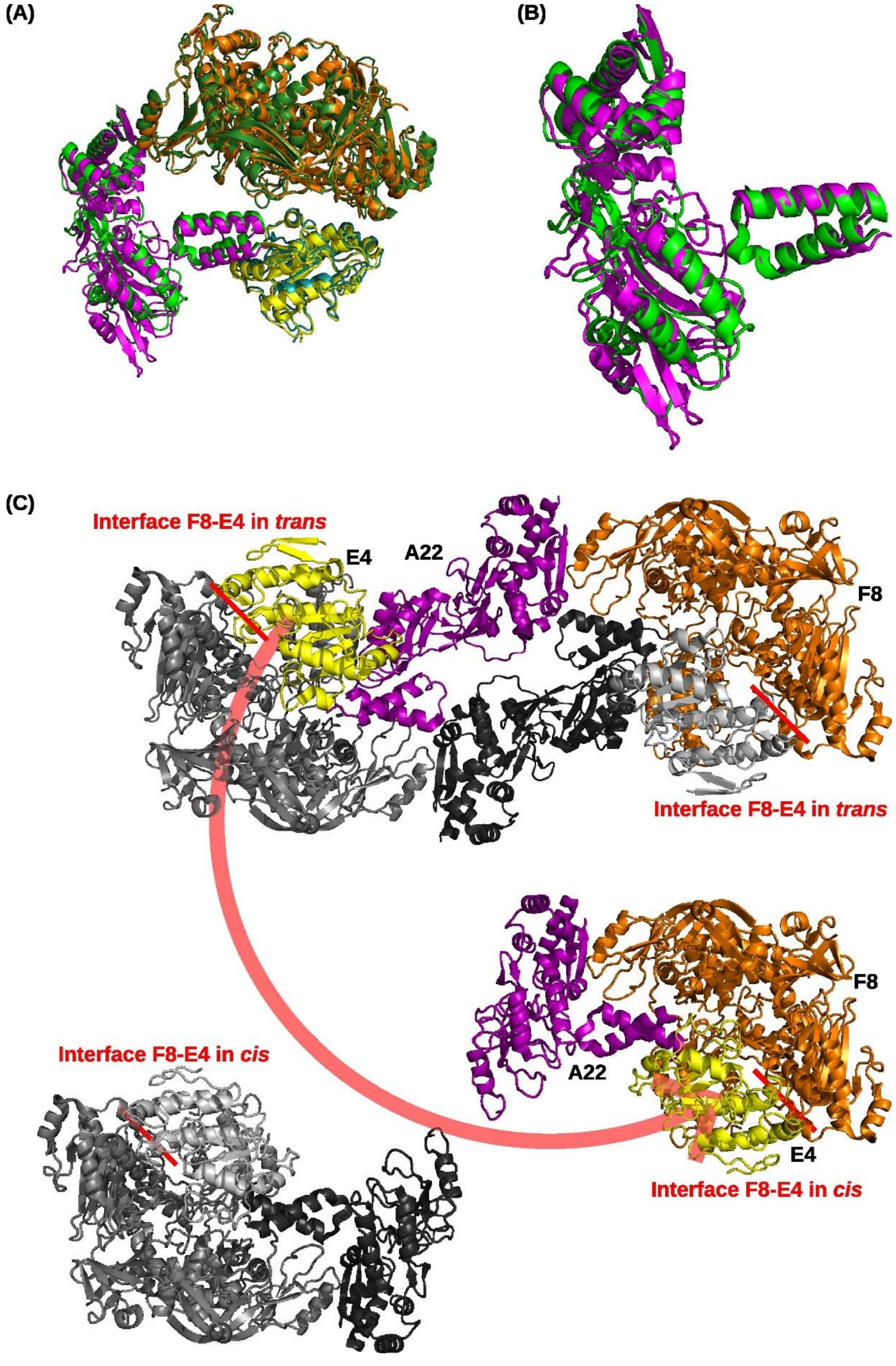

**S3 Fig. Comparison of the holoenzyme heterotrimer of VACV polymerase with the one of MPXV.** (**A**) Superposition of the VACV holoenzyme (orange, violet, yellow) and the MPXV one (pdb entry 8hm0, dark green, light green, cyan) based on E9 and D4. (**B**) Superposition of A22 from MPXV virus (green) onto A20 of VACV (magenta). (**C**) Top: interpretation of the hexamer structure (pdb entry 8hlz) as a dimer of extended trimers. One subunit is colored using our standard color scheme, the other one in colored in shades of gray. The F8-D4 interface is formed in trans. Bottom: Formation of the compact monomer in pdb entry 8hm0 by the dissociation and rebinding of D4 forming now the F8-E4 interface in cis.

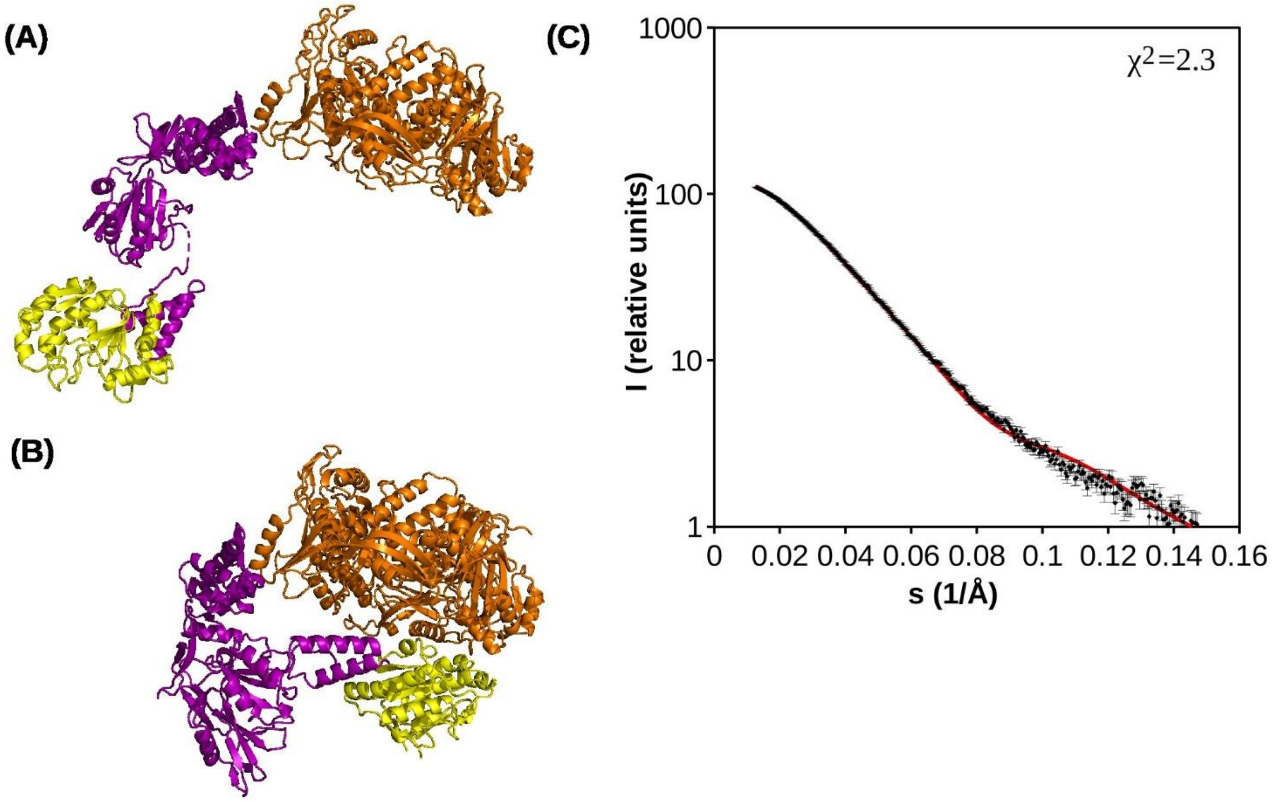

**S4 Fig. Explanation of the SAXS curve of VACV holoenzyme**. (**A**) Extended form of the VACV heterotrimer based on pdb entry 8hlz. (**B**) Compact heterotrimer of the VACV polymerase holoenzyme (**C**) SAXS curve of the VACV holoenzyme (as in Fig 3) together with a calculated curve (red) based on a mix of 58 % of an extended VACV E9-A20-D4 holoenzyme (A) and 42 % of the compact form (B).
